# Comparison of three glycoproteomic methods for the analysis of CHO cells treated with 1,3,4-O-Bu_3_ManNAc

**DOI:** 10.1101/2020.02.18.954198

**Authors:** Joseph L. Mertz, Shisheng Sun, Bojiao Yin, Michael J. Betenbaugh, Kevin J. Yarema, Hui Zhang

## Abstract

Comprehensive analysis of the glycoproteome is critical due to the widespread importance of this post-translational modification to protein function, and difficult because of the tremendous complexity it exhibits. Here we compared three glycoproteomic analysis methods, a recently described chemoenzymatic glycoproteome analysis methods, N-linked glycans and glycosite containing peptides (NGAG), Solid-phase extraction of N-linked glycoproteins (SPEG), and hydrophilic interaction liquid chromatography (HILIC), for the analysis of N-linked glycosites of Chinese hamster ovarian (CHO) cells treated with 1,3,4-O-Bu_3_ManNAc. The NGAG protocol resulted in substantially increased glycosite identifications over both SPEG and HILIC. Interestingly, while the glycosites identified by SPEG and HILIC overlapped strongly, NGAG identified many glycosites not observed in either of the other two methods. Further, utilizing the enhanced intact glycopeptide identification afforded by the NGAG workflow, we also found that of the sugar analog 1,3,4-O-Bu_3_ManNAc increases sialylation of proteins secreted by CHO cells, including an ectopically expressed human proteins.

## Introduction

Glycosylation is a widespread protein modification that plays a role in protein folding, stability, protein−ligand interactions, biological function, and pathogenesis (1). Because each glycoprotein can exhibit multiple glycosylation sites (glycosites) and each glycosite can exhibit a number of glycan structures, glycosylation is also the most complex modification, with the potential for a single protein isoform to exhibit thousands of glycosylation combinations (2). Further, specific glycan structures at specific glycosites have been associated with altered function and disease (3), and thus comprehensive systematic analysis is both challenging and crucial.

Virtually all therapeutic proteins are glycosylated – with notable exceptions such as insulin and human growth hormone – and glycosylation can influence the conformation, stability, pharmacokinetic profile, *in vivo* activity, and immunogenicity of recombinant proteins (4). One significant and well-documented contribution of glycosylation to the pharmacokinetic profile – generally encompassing absorption, distribution, metabolism and excretion (ADME) – is increased half-life in circulation, which greatly affects dosing strategies and thus ultimate success of a drug (5,6). Terminal sialic acid residues, in particular, have been shown to increase circulatory half-life by masking galactose residues from Galactose-specific receptors in the liver (7) and other would-be terminal glycan residues from lectin-like receptors elsewhere in the body (8,9).

Mammalian cell expression systems predominate therapeutic protein production and among these Chinese hamster ovary (CHO) cells are by far the most prevalent. This is due to their capacity for high-density culture in serum-free conditions, a vast accumulation of expertise surrounding these cells, the development of efficient gene introduction techniques, and glycosylation profiles that are compatible for use in humans (10). While the glycosylation machinery in CHO cells is highly similar to humans, differences do exist - notably they lack an α-2,6-sialyltransferase ortholog, encoded by ST6GAL1 in humans, which decreases their overall level of sialylation (11–13). Thus, strategies have been introduced to glycoengineer CHO cell culture systems to optimize glycosylation patterns both genetically and metabolically. Genetic approaches include exogenous expression of glycosylation machinery genes ST6Gal1, GNTIV, and GNTV while metabolic approaches involve additives to culture media to help drive desired forms of glycosylation (14). One such metabolic additive is 1,3,4-O-Bu_3_ManNAc, a synthetic analogue of the sialic acid precursor N-acetyl-D-mannosamine (ManNAc) with the addition of the short chain fatty acid butyrate at the 1, 3, and 4 positions to increase uptake across the cell membrane. This analogue strongly increases metabolic flux into the sialylation pathway, exhibiting increased sialic acid overall as measured by periodate resorcinol assay (15) as well as increased sialylation of specific proteins such as epidermal growth factor receptor (EGFR) (16) and erythropoietin (EPO) (17) with low toxicity.

Glycoproteomics research has progressed rapidly over the last decade, propelled by advances in sample preparation techniques, mass spectrometry (MS) based technologies, and data processing. Innovations in affinity-based capture of glycans, glycopeptides, and glycoproteins by lectins, antibodies, or chemical affinity resins (18,19) have increased coverage of the glycoproteome (20–22). Solid-phase extraction of N-linked glycoproteins (SPEG), which utilizes covalent bonds between hydrazide beads and oxidized glycan moieties on glycosite-containing peptides followed by release by PNGase was one of the first examples of such a workflow, and it has seen continued use since its introduction (22–24). Hydrophilic interaction liquid chromatography (HILIC), is a collective term for purification techniques that generally utilize solid phases with neutral polar moieties to retain an aqueous layer, and thereby sequester glycopeptides via their hydrophilic glycan moieties from a less polar mobile phase (25). Subsequent PNGase treatment and LC-MS/MS allows proteome-wide glycosite identification as in SPEG workflows.

Glycosite identification has recently been succeeded on the cutting edge of glycoproteomics research by intact glycopeptide analysis, which has been made feasible by the introduction of higher-energy collisional dissociation (HCD) with glycan-originating oxonium ions alongside *b* and *y* ions from the peptide backbone (26). The increase in search space brought by various glycan chains makes accurate identification difficult and has led to various strategies to restrict or guide the peptide sequence matching process for intact glycopeptides. A particularly powerful method known as solid phase extraction of N-linked glycans and glycosite-containing peptides (NGAG) (27) combines chemical and enzymatic modifications of peptides to allow covalent binding to a solid phase and specific release of N-glycosylated peptides. Briefly, after trypsin digestion: glycopeptides are guanidinated to block lysine ε-amino groups; covalently bound to an aldehyde solid phase via the terminal primary amine; treated with aniline to modify carboxyl groups – including, importantly, aspartic acid and sialic acid residues; deglycosylated and glycosylated asparagines deaminated to aspartic acid by PNGaseF treatment; peptides containing aspartic acid residues newly formed by deglycosylation (and not converted by the previous aniline treatment) are selectively released by Asp-N; released peptides are analyzed by LC-MS/MS to identify glycosites; and finally these glycosites are used for improved peptide sequence match (PSM) assignment of parallel intact glycopeptide samples. Data analysis in the NGAG workflow is performed by the GPQuest software package (28,29), which integrates perfectly with the NGAG protocol to match intact glycopeptide LC-MS/MS spectra to glycan and peptide databases generated from the same samples. Though, notably, the utility of GPQuest extends to other glycoproteomics protocols as well.

To test the potential of shifting the sialylation equilibrium toward more therapeutically efficacious states, we treated CHO-K1 cells expressing recombinant human erythropoietin (EPO) with the ManNAc analogue 1,3,4-O-Bu_3_ManNAc and analyzed them by the state-of-the-art and highly comprehensive NGAG technique recently developed by our group. Further, to characterize glycoproteome detection afforded by the NGAG pipeline, we also performed the more conventional HILIC and SPEG analyses in parallel. Our results show 1,3,4-O-Bu_3_ManNAc to be a potent glycoengineering tool for increasing sialylation throughout the proteome – including the endogenously expressed EPO – and that NGAG produces substantially more glycopeptide identifications with a surprising degree of orthogonality to either HILIC or SPEG.

## Methods

### CHO cell culture and protein harvest

Two biological replicates of CHO cells ectopically expressing human EPO protein were cultured in suspension with and without 1,3,4-O-Bu_3_ManNAc treatment. The cells were pelleted and washed twice with PBS while the protein in the medium was concentrated and buffer exchanged by 10 KD filter. Protein was harvested from each sample with 10 mL 8 M urea in 1 M NH_4_HCO_3_ followed by two rounds of sonication with 40-60 cycles per round. Protein was reduced by 10 mM TCEP (Thermo Fisher Scientific), alkylated with 16.5 mM iodoacetamide (Sigma-Adrich), and digested with sequencing-grade trypsin (Promega) at 1:50 enzyme:substrate ratio. Peptides were desalted using a C18 solid phase extraction (SPE) cartridge (Waters) and dried down by SpeedVac.

### SPEG enrichment of deglycosylated peptides

Cis-diols of glycan moieties on tryptic glycopeptides were oxidized by 10 mM NaIO_4_ for 1 hr in the dark and the oxidized glycopeptides then purified by C18 column. 1% aniline was added to the oxidized glycopeptides, which were conjugated to 100 μl hydrazide beads per 1 mg peptides overnight at pH<6. The beads were washed three times each with 50% ACN, 1.5 M NaCl, H_2_O and 25 mM NH_4_HCO_3_ with 20 s vortexing at each wash. Peptides were released from their glycans and the beads by PNGase F cleavage in 25 mM NH_4_HCO_3_. The supernatant plus one wash of the beads with H_2_O and one with 50% ACN was collected for analysis of the deglycosylated and deamidated peptides for SPEG glycosite analysis.

### HILIC enrichment of intact glycopeptides, deglycosylated peptides, and glycans

Tryptic glycopeptides from each sample were enriched using a HILIC column (SeQuant, Southborough, MA). After desalting, peptides were reconstituted in a final solvent composition of 80% ACN/0.1% TFA. The HILIC column was conditioned twice with 0.1% TFA and twice with 80% ACN/0.1% TFA, the sample was loaded and washed three times with 80% ACN/0.1% TFA, then bound glycopeptides were eluted in 0.1% TFA. The samples were dried via SpeedVac and resuspended in 0.2% FA if they were to be used for intact glycopeptide LC-MS/MS analysis or 25 mM NH_4_HCO_3_ for separate glycosite and glycan analysis. For glycosite and glycan analysis, N-glycans and peptides were separated by PNGase F digestion overnight. Deglycosylated and deamidated peptides were purified by C18 column and resuspended in 0.2% FA for LC-MS/MS. For N-glycan analysis, the glycans were purified using HyperSep Hypercarb SPE cartridges (Thermo Fisher Scientific). The columns were conditioned with 100% acetonitrile and 1% TFA, then the sampled loaded in 0.1% FA, washed with 0.1% FA, and eluted in 80% ACN/0.1% FA.

### NGAG enrichment of deglycosylated peptides and glycans

Lysines on tryptic glycopeptides from each sample were guanidinated using 1.425 M NH_3_.H_2_O and 0.6 M o-methylisourea for 30 min at 65 °C then purified by C18 column. Guanidinated peptides were conjugated by their amino terminals to 500 mL AminoLink beads per 1 mg peptide overnight at RT. Carboxyl groups – on both glycans and the peptide backbones – were modified by aniline in the presence of 0.5 M 1-ethyl-3-(3-dimethylaminopropyl)carbodiimide (EDC) in 200 mM MES buffer, pH 5, at RT overnight. Peptides with non-modified aspartate residues were removed by Asp-N at 37 °C overnight, then N-glycans released and peptides deaminated by overnight digestion with PNGase F. Deglycosylated peptides were released at their nascent aspartate residues by overnight digestion with Asp-N in 25 mM NH_4_HCO_3_ and the collected in the supernatant and prepared for LC-MS/MS analysis.

### Mass spectrometry analysis of glycans

N-glycans released by the NGAG protocol and purified on HyperSep cartridges were dried, then resuspended in deionized water. Samples of this resuspension were spotted onto a MALDI plate along with equal volume 2,5-dihydroxybenzoic acid/N,N-dimethylaniline (DHB/DMA) matrix containing 100 mg/ml DHB, 2% DMA in 50% acetonitrile and 0.1 mM NaCl. Analysis was performed on an Axima MALDI Resonance mass spectrometer (Shimadzu) with the laser power set to 100 for 200 total shots per spot. Glycan structures were elucidated from the most abundant peaks in the 800-2000 Da and 2000-3000 Da spectra by Glyco-Peakfinder (EuroCarbDB).

### LC-MS/MS analysis

C18-purified tryptic peptides, and deglycosylated peptide samples from SPEG and HILIC were analyzed on a Velos Pro mass spectrometer while HILIC-purified intact glycopeptides and NGAG deglycosylated peptide samples were analyzed on a Q-Exactive mass spectrometer (Thermo Fisher Scientific, Bremen, Germany). For the Q-Exactive analyses, peptides were separated on a Dionex Ultimate 3000 RSLC nano system with a 75 μm × 15 cm Acclaim PepMap100 separating column (Thermo Scientific). Buffer A consisted of 0.1% formic acid and Buffer B 0.1% formic acid 95% acetonitrile at 250-290 nL/min. For NGAG glycosite analysis deglycosylated peptides were separated by a gradient profile of 4-10% Buffer B over 6 minutes, 10-28% B over 84 minutes, 28-35% B over 10 minutes, 35-90% B over 10 minutes, 90% B for 8 minutes, 90-4% B over 2 minutes, then 10 minutes at 4%. The intact glycopeptide analysis utilized a slightly steeper gradient profile with 4% Buffer B for 5 minutes, 4-25% B over 95 minutes, 25-95% B over 5 minutes, 95% for 10 minutes, 95-45 over 1 minute, at 4 minutes at 4%. MS/MS on these samples was performed with 1.8-2.0 kV electrospray voltage. MS1 spectra were collected in the Orbitrap with AGC target of 3×10^6^ and 70,000 resolution from 400 to 2,000 m/z. MS2 spectra were collected using HCD fragmentation of the top 9-15 most abundant ions at 17,500 resolution and 100 m/z fixed first mass. A dynamic exclusion of 15 seconds was used for intact analysis and 30 seconds for NGAG glycosite analysis. SPEG, HILIC, and Global experiments were analyzed on an Orbitrap Velos Pro instrument. Peptides were separated on an UltiMate UPLC system with a 75 μm × 15 cm Acclaim PepMap100 separating column (Thermo Scientific) at 300 nL/min. Separation was achieved by a gradient profile of 4-8% Buffer B over 6 minutes, 8-35% B over 84 minutes, 35-45% B over 10 minutes, 45-90% over 10 minutes, and 10 minutes at 90%. Electrospray voltage was 2.2 kV. MS1 spectra were collected from 400 to 1,800 m/z with a resolution of 30,000. The top 10 most abundant ions with a 25 s dynamic exclusion were fragmented by HCD and MS2 were collected at 7,500 resolution with 100 m/z fixed first mass.

### Global and glycosite data analysis

LC-MS/MS data were searched by MaxQuant (v1.5.8.3) against the Uniprot Cricetulus griseus proteome database downloaded May 5^th^, 2016. Global proteome data was searched using up to two missed cleavages, 20 ppm peptide mass tolerance for the first search and 4.5 ppm for the main search, carbamidomethylation of C (+57.0215 Da) as a static modification, oxidation of M (+15.9949 Da) and acetylation of protein N-terminal (+42.0106 Da) as dynamic modifications, five modifications per peptide, a minimum peptide length of seven amino acids, and 1% FDR against a reversed decoy proteome database. For SPEG and HILIC glycosite data searches, deamination of N (+0.9840) was added as a dynamic modification. NGAG glycosite data searches used a custom proteome created by replacing the N of all potential N-glycosylation sites (N-X-S/T motifs, X≠P) with the 168.9642 Da dummy amino acid “U.” NGAG searches also used a custom enzyme “Trypsin + U” created on the MaxQuant enzyme list, with cleavage at the N-terminal side of “U” in addition to trypsin K and R cleavage sites. NGAG search parameters included up to two missed cleavages for “trypsin+U” digestion; 20 ppm mass tolerance for first search and 6 ppm for main search; carbamidomethylation of C (+57.0215 Da) as a static modification; dynamic modifications of: U→D (−53.9373 Da), U not at N termini→N (−54.9213 Da), guanidination of K (+42.0218 Da), aniline at protein C termini, D and E not at N termini, and K and R at any C terminus (+75.0473 Da), two anilines on D and E at protein C termini (+150.0946 Da), guanidination and aniline on K at any C termini (+117.0691 Da) and oxidation of M (+15.9949 Da); six modifications were allowed per peptide, a minimum peptide length of 7 amino acids, and a 1% FDR versus a reversed decoy proteome database.

### Intact glycopeptide data analysis

Intact LC-MS/MS data were analyzed by the GPQuest software package developed in-house (29). Briefly, glycopeptide MS/MS spectra were extracted by assaying the five most abundant MS2 ions for oxonium ions of HexNAc (204.087 Da) ion and one other glycan (138.055 Da, 163.061 Da, 168.066 Da, 274.093 Da, 292.103 Da and 366.140 Da) with 50 ppm tolerance. These spectra were matched with a 10 ppm mass error to the candidate database comprised of glycans identified by PNGase F release and MALDI analysis and glycosite-containing peptides identified by the combined glycosite analyses.

## Results

### Analysis of CHO cells with three glycoproteomic approaches to identify changes of glycosylation sites with sugar analog treatment

To determine how NGAG fits into the greater framework of glycopeptide enrichment methodologies we performed a comparison of the glycopeptide enrichment obtained with two widely used approaches, SPEG and HILIC, to the NGAG approach (summarized in Figure 1). SPEG and HILIC utilize orthogonal enrichment mechanisms: SPEG covalently binds glycan carbonyl groups (Fig. 2A), while HILIC sequesters the more hydrophilic glycopeptides from the rest of the tryptic proteome (Fig. 2B). NGAG uses yet another mechanism to enrich for glycopeptides in which it binds all peptides by the amino terminal but only releases those exhibiting N-glycosylation after a targeted chemoenzymatic modification process (Fig. 2C).

**Figure 1.**
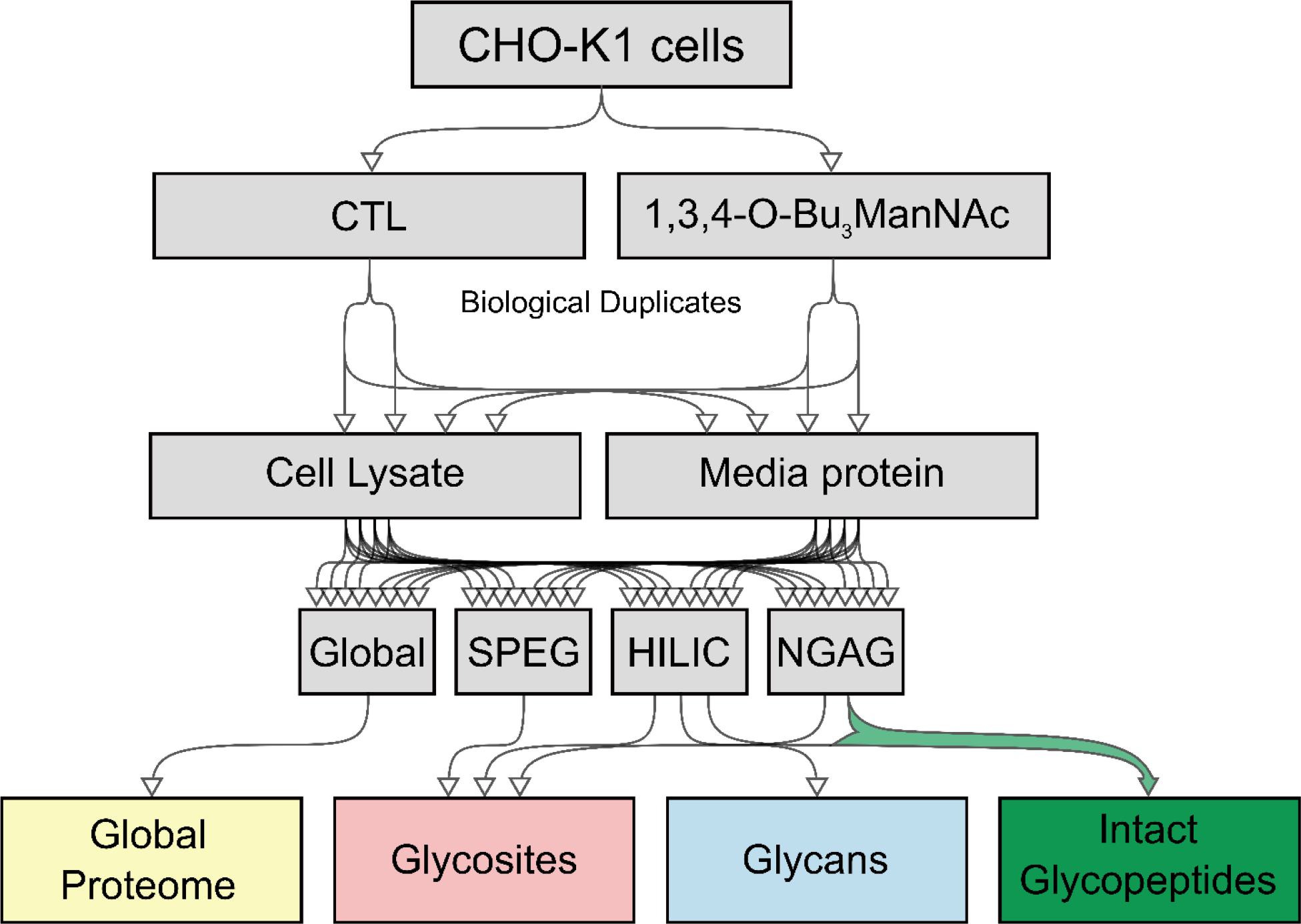
Experimental workflow. Duplicate samples of CHO-KI cells expressing endogenous human erythropoietin were untreated or treated with 1,3,4-O-Bu_3_ManNAc. Protein was harvested and digested by trypsin, the global proteome was analyzed, and equal amounts were used for each glycoproteome workflow to generate glycosite, glycan, and intact glycopeptide data.

**Figure 2.**
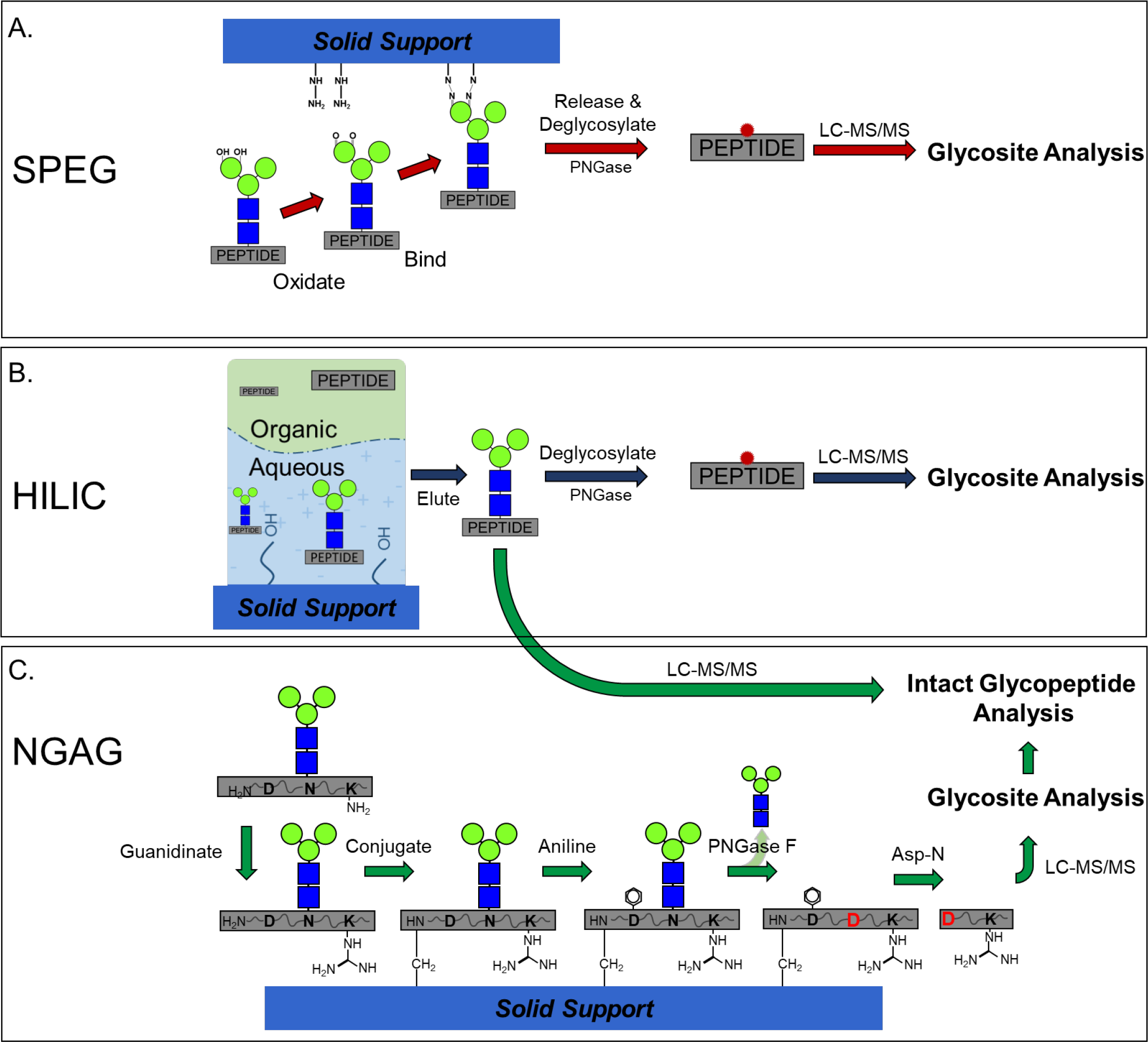
Schematic of each glycoproteome workflow. A) In SPEG glycosite identification analysis, glycopeptides are captured by covalent binding of oxidized glycan moieties to hydrazide beads followed by release by PNGase. B) In HILIC glycosite analysis, glycopeptides are sequestered in the aqueous phase retained by the polar moieties of the column, eluted, then deglycosylated by PNGase. C) In NGAG intact glycopeptide analysis, a stepwise chemoenzymatic process extracts glycopeptides via the peptide N-terminus, deglycosylated by PNGase, and ultimately released by Asp-N cleavage at the nascent aspartic acid residues for glycosite analysis. The resulting peptide library is combined with intact glycopeptide enrichment via HILIC – without PNGase treatment – for more efficacious intact glycopeptide identification.

**Figure 3.**
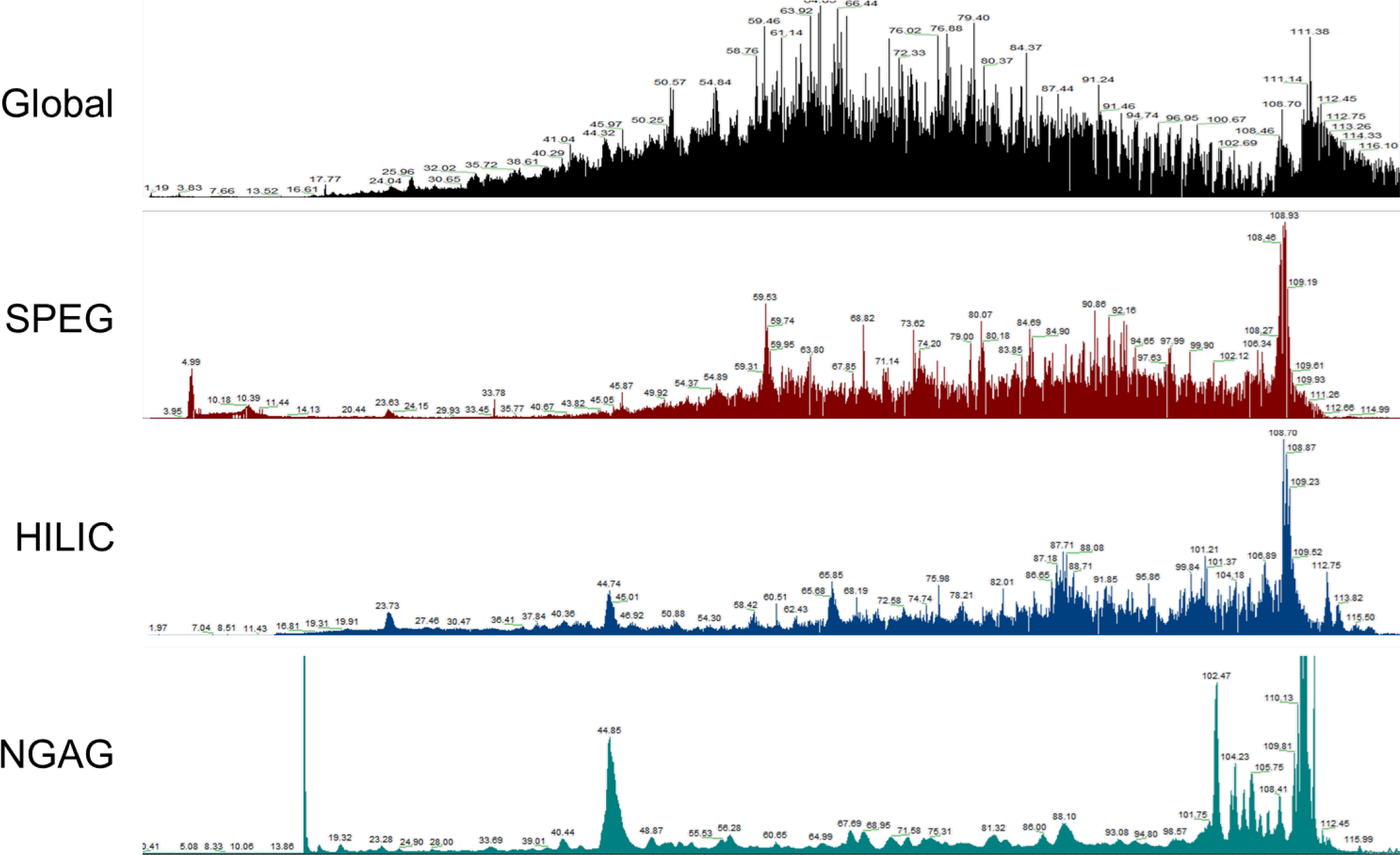
Gross analysis of the LC-MS spectra suggest similarity between SPEG and HILIC preparations, with a more distinct trace from NGAG.

We performed these three approaches in parallel on matching starting quantities of the same secreted protein samples from CHO cell cultures. We centrifuged the culture mixture and collected the supernatant for analysis of the secreted proteomes and the cell pellet for cell proteomes). After protein extraction, trypsinization, and desalting, we used the resulting tryptic peptides from each condition for the various proteome enrichment methods. After each enrichment workflow, we analyzed each by triplicate single-shot LS-MS/MS runs.

First, comparing total glycosite identifications from cell culture media we see similar total numbers of identifications, with NGAG returning the most unique sites, 586, HILIC with 477, and SPEG with 301. The performance of NGAG represents a 23% increase in identifications over HILIC and a 94% increase over SPEG. When examining the overlap between the three methods (Fig. 4), we saw high levels of glycosite overlap between SPEG and HILIC – 70.1% of those from SPEG were also identified by HILIC, 44.2% of HILIC also identified by SPEG, and 37.2% of the 567 identified between the two were identified by both. Interestingly, the glycosites identified by NGAG exhibit much less similarity to either of the other two methods, nor indeed both combined. NGAG identified 126 glycosites in common with SPEG – 21.5% of the total NGAG glycosites and 41.9% of the SPEG total. NGAG and HILIC, meanwhile, overlapped on 29.5 of NGAG glycosites and 36.3% of HILIC. Of the 953 glycosites identified altogether, only 99 or 10.4% were identified by all three methods and 386 or 40.5% were identified only by NGAG, a number with greatly surpasses the 63 sites (6.6%) unique to SPEG or the 192 (20.1%) unique to HILIC. When taken together, the higher number of identifications – 586 by NGAG topping even the 567 combined between SPEG and HILIC – and the notable lack of overlap suggest NGAG represents an enrichment methodology with increased performance that may extract a portion of the glycoproteome not commonly studied up to this point. MALDI-TOF glycan peaks matched to glycan structures by GlycoPeak finder identified 60 unique glycans, with high amounts of overlap between conditions (Fig. 5). These were used for downstream analysis of intact glycopeptides below.

**Figure 4.**
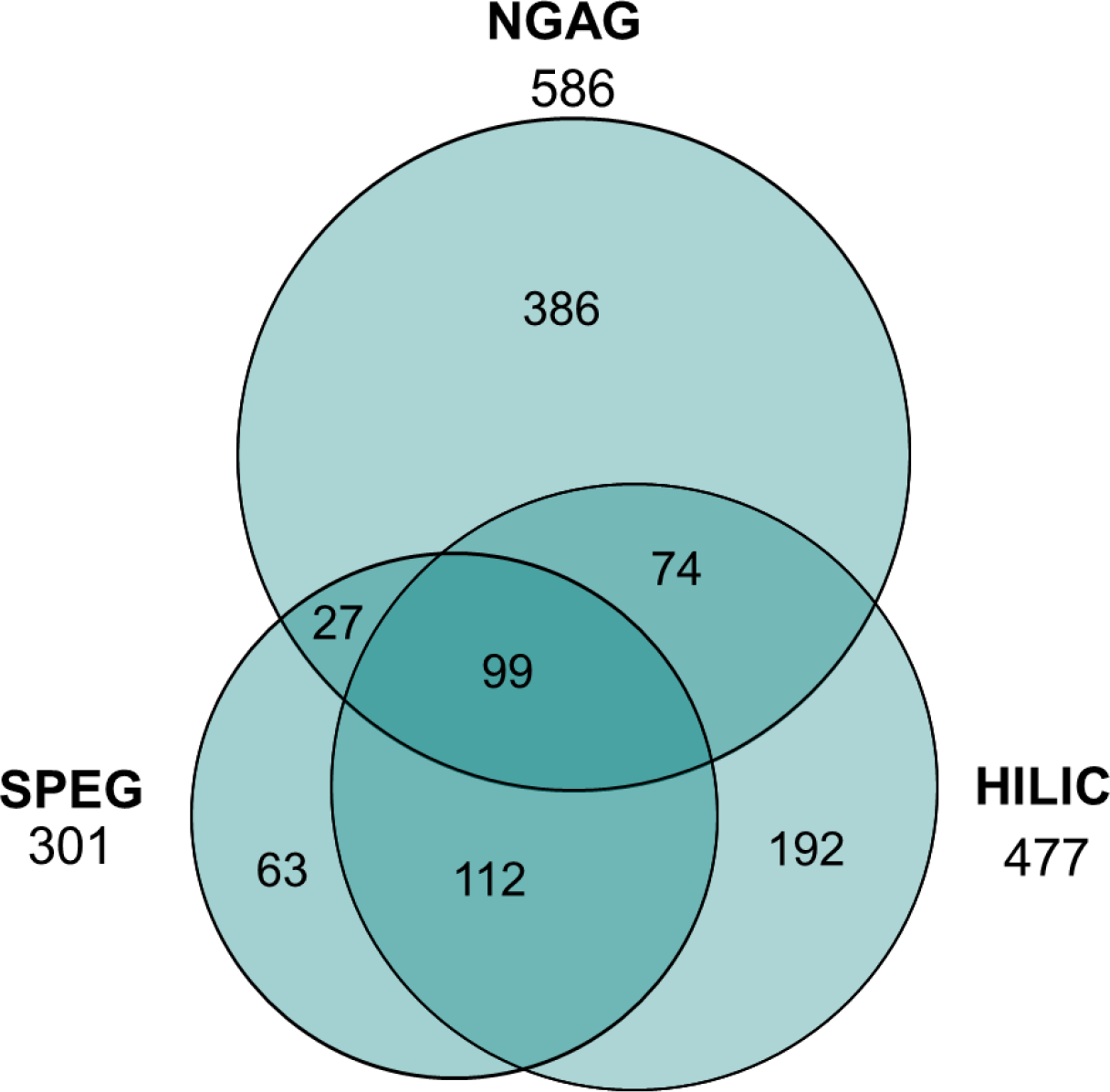
SPEG and HILIC exhibit strong overlap in glycosite identification with NGAG exhibiting orthogonality. Overall, NGAG identified the largest number of glycosites, 586, from media samples with HILIC identifying 477 and SPEG 301. Interestingly, while SPEG and HILIC overlap significantly, the vast majority of glycosites identified by NGAG were not observed in the other two datasets.

**Figure 5.**
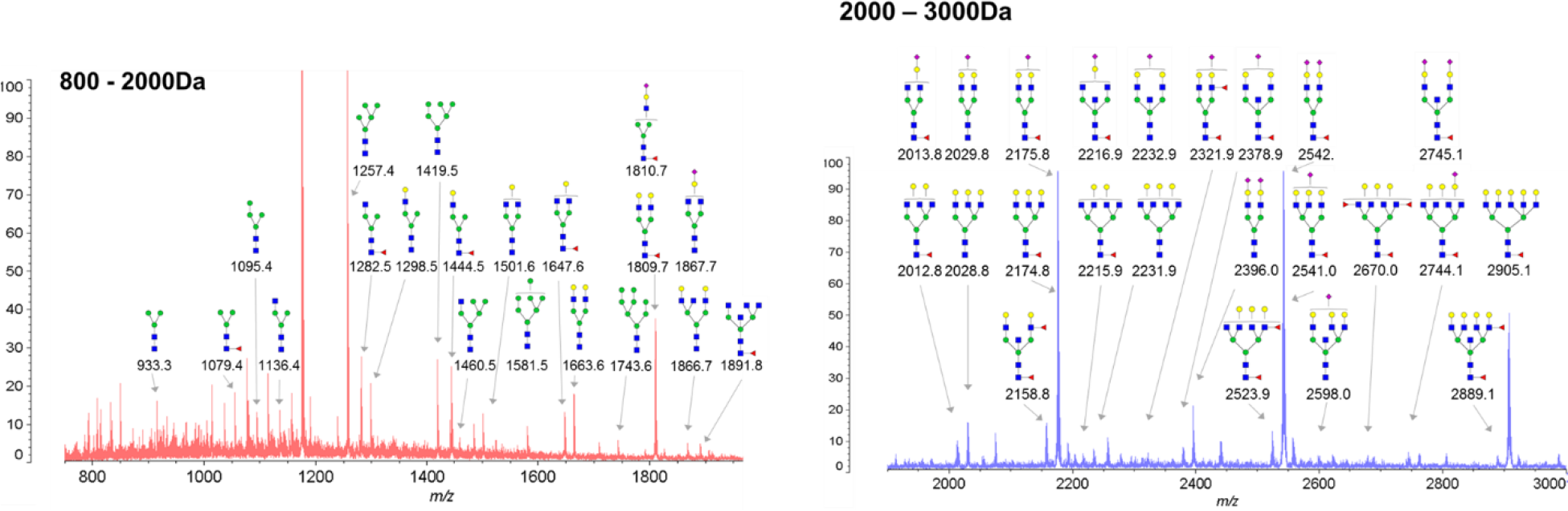
Glycan analysis by MALDI-TOF identified 60 unique glycans across the range of samples

The SPEG and HILIC pipelines were also performed on protein harvested directly from the CHO cells, which allowed comparison of intracellular and extracellular glycosylation patterns. Overall, glycosites identified from cellular proteins differed vastly from their secreted counterparts, a result borne out by both enrichment methods. The number of unique glycosites identified from cellular proteins was much higher for both enrichment methods, with 729 cell vs 301 media in SPEG and 721 cellular vs 477 media in HILIC. Further, the glycosylation patterns of individual proteins differed between intra and extracellular localization: numerous proteins identified by each method exhibited glycosylation at a given site in all cellular fractions but in no extracellular fractions (or conversely, exhibited glycosylation in all extracellular fractions but no intracellular fractions), while other glycosites on the same protein were observed in both intra- and extracellular fractions. These data suggest that these glycosites play a role in the secretion of these proteins – or that intra- vs extracellular location plays a role in their glycosylation.

### Quantitative analysis of N-glycosites using three glycoproteomic methods reveals similarity and complementary of three methods

Using precursor ion intensity-based label free quantification, we were able to quantify relative abundance of glycosite containing peptides between conditions and enrichment strategies. Overall, glycosite quantitation between the three methods exhibit reasonable correlation, though agreement between rank-order abundances of glycosites was much higher between different fractions of the same enrichment method than between different enrichments of the same material (Fig. 6). Thus, the biases in glycopeptide capture efficacy between these methods can outweigh biological sources of abundance differences, which is not unexpected but warrants caution, particularly when comparing between glycosite abundances.

**Figure 6.**
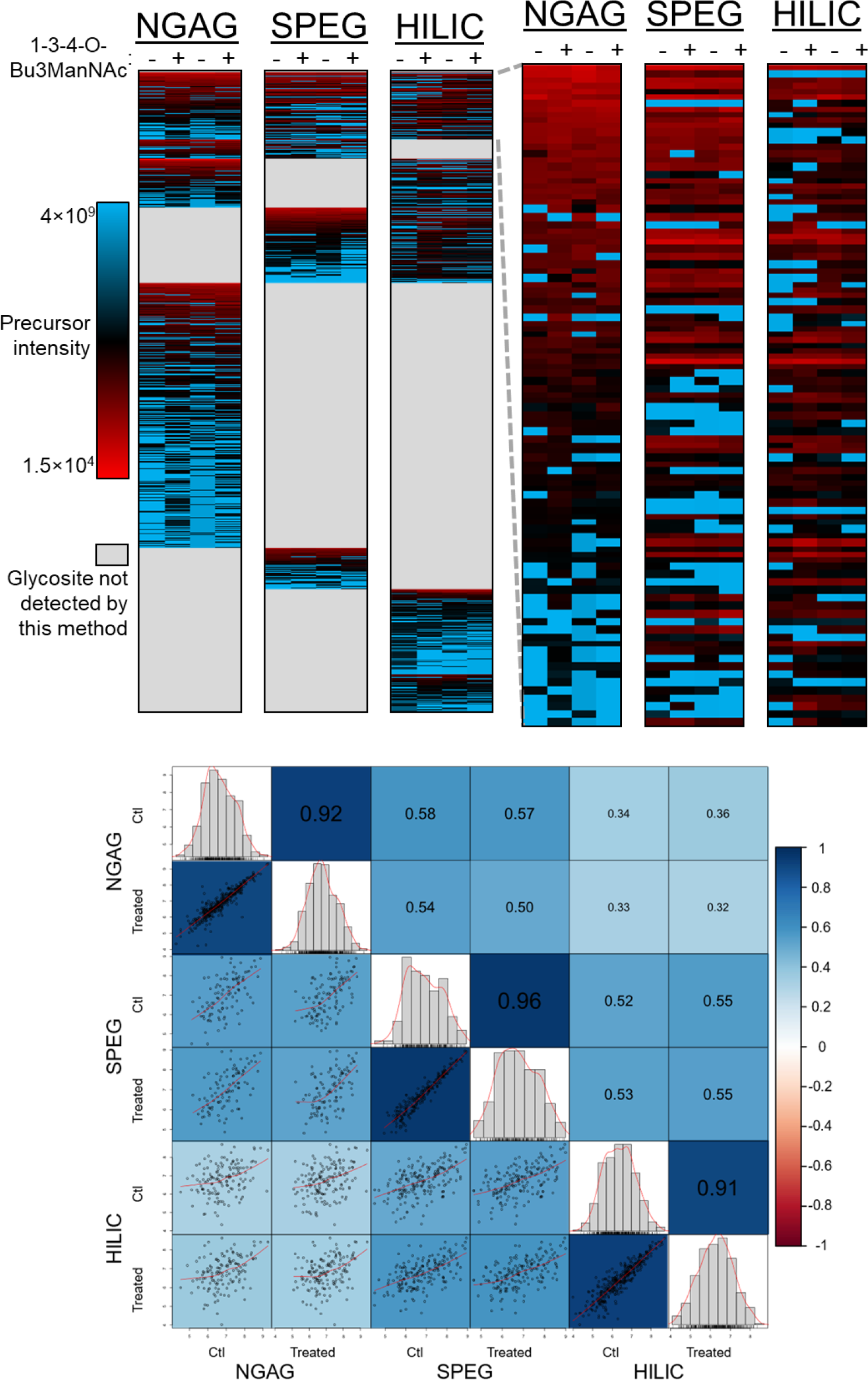
Glycosite quantification by precursor intensity reveals reasonable correlation between the different approaches in the glycosites identified by multiple approaches. Overall, much higher correlation was observed between the results of the same workflow from different treatment samples, with notably lower correlation between NGAG and HILIC results.

### Intact glycopeptide analysis reveals glycosylation heterogeneity at different glycosylation sites

We sought to examine glycan-at-glycosite heterogeneity in the CHO cell secreted proteome and how priming sialylation machinery may alter this by comparing the secreted proteomes of CHO cells with and without 1-3-4-O-Bu3ManNAc treatment. This N-Acetylmannosamine ester efficiently crosses cellular membranes and enters the sialylation pathway as a committed precursor to sialic acid, thereby potentially driving inclusion of sialic residues during glycosylation. For analysis to place glycans at a given glycosite, intact glycopeptides are required, so we employed the NGAG workflow using HILIC-based purification without PNGase treatment and LC-MS/MS analysis of the resulting intact glycopeptides. Using GPQUEST, a software suite developed in house, these data were then searched against a glycosite-only peptide database generated from the NGAG glycosite analyses and an N-glycan database generated from the released N-glycan MALDI-TOF data to identify and quantify intact glycopeptides. Narrowing the peptide and glycan databases down from the entire proteome in this way greatly reduces search space and allows much more accurate peptide + modifying glycan PSM assignment in much less time. By this method, in total from all 4 media fractions we identified 686 unique intact glycopeptides, which represented combinations of 55 unique glycans at 193 unique glycosites from 124 different proteins (Summarized in Table 1).

**Table 1.**
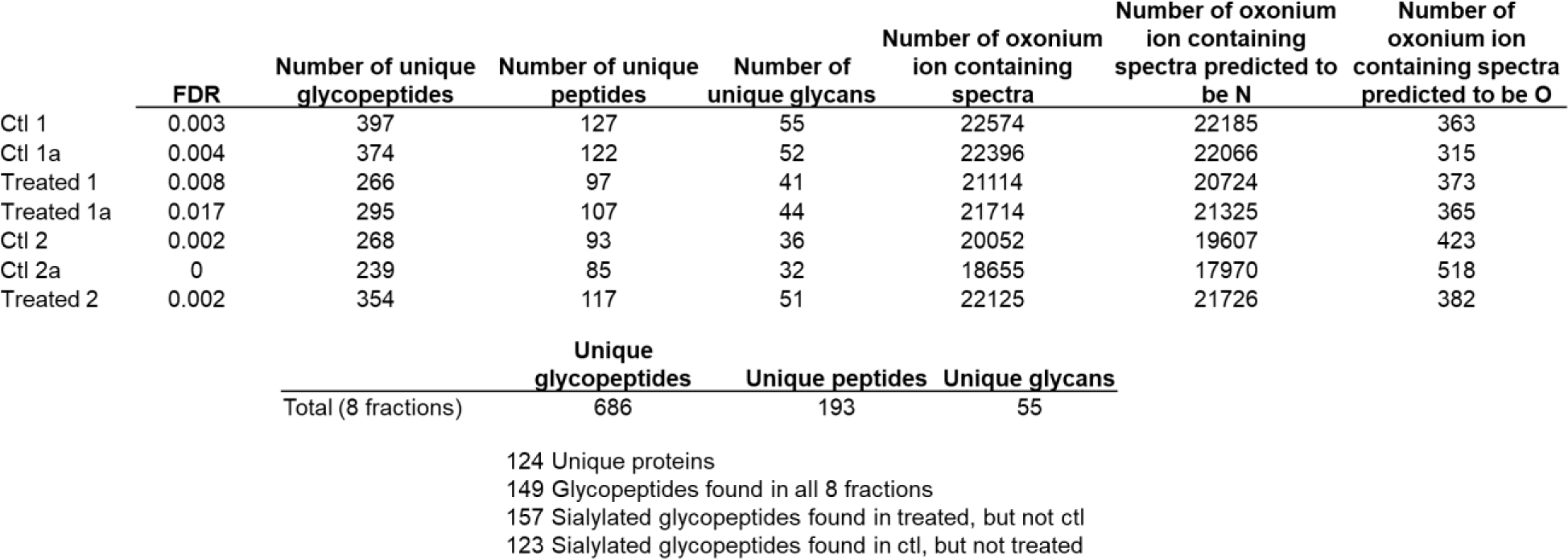
NGAG intact glycopeptide analysis of media preparations identified 686 unique glycopeptides resulting from combinations of 193 unique peptides and 55 unique glycans. Notably, 1,3,4-O-Bu_3_ManNAc did not cause a significant increase in glycopeptide or glycan number.

|  | **FDR** | **Number of unique glycopeptides** | **Number of unique peptides** | **Number of unique glycans** | **Number of oxonium ion containing spectra** | **Number of oxonium ion containing spectra predicted to be N** | **Number of oxonium ion containing spectra predicted to be O** |
| --- | --- | --- | --- | --- | --- | --- | --- |
| Ctl 1 | 0.003 | 397 | 127 | 55 | 22574 | 22185 | 363 |
| Ctl 1a | 0.004 | 374 | 122 | 52 | 22396 | 22066 | 315 |
| Treated 1 | 0.008 | 266 | 97 | 41 | 21114 | 20724 | 373 |
| Treated 1a | 0.017 | 295 | 107 | 44 | 21714 | 21325 | 365 |
| Ctl 2 | 0.002 | 268 | 93 | 36 | 20052 | 19607 | 423 |
| Ctl 2a | 0 | 239 | 85 | 32 | 18655 | 17970 | 518 |
| Treated 2 | 0.002 | 354 | 117 | 51 | 22125 | 21726 | 382 |

|  | **Unique glycopeptides** | **Unique peptides** | **Unique glycans** |
| --- | --- | --- | --- |
| Total (8 fractions) | 686 | 193 | 55 |

124 Unique proteins
149 Glycopeptides found in all 8 fractions
157 Sialylated glycopeptides found in treated, but not ctl
123 Sialylated glycopeptides found in ctl, but not treated

### Intact glycopeptide analysis revealed increased numbers of glycosites with multiple sialic acids after treatment of sugar analog

Comparison of intact glycopeptide data from 1-3-4-O-Bu3ManNAc-treated and control CHO cells revealed that overall glycopeptide numbers were not significantly altered by 1-3-4-O-Bu_3_ManNAc treatment. However, when focusing on glycan composition, we observed significantly increased inclusion of sialic acid residues (Fig. 7A). This was observable in two ways: the proportion of glycopeptides with 2, 3, and 4 sialic residues increased upon 1-3-4-O-Bu_3_ManNAc treatment, while the proportion of glycopeptides with 0 or 1 sialic residues was decreased. This zero-sum pattern in the data suggests 1-3-4-O-Bu3ManNAc did not increase overall glycosylation in any significant way but did increase the likelihood that a glycan would be sialylated (Fig. 7B).

**Figure 7.**
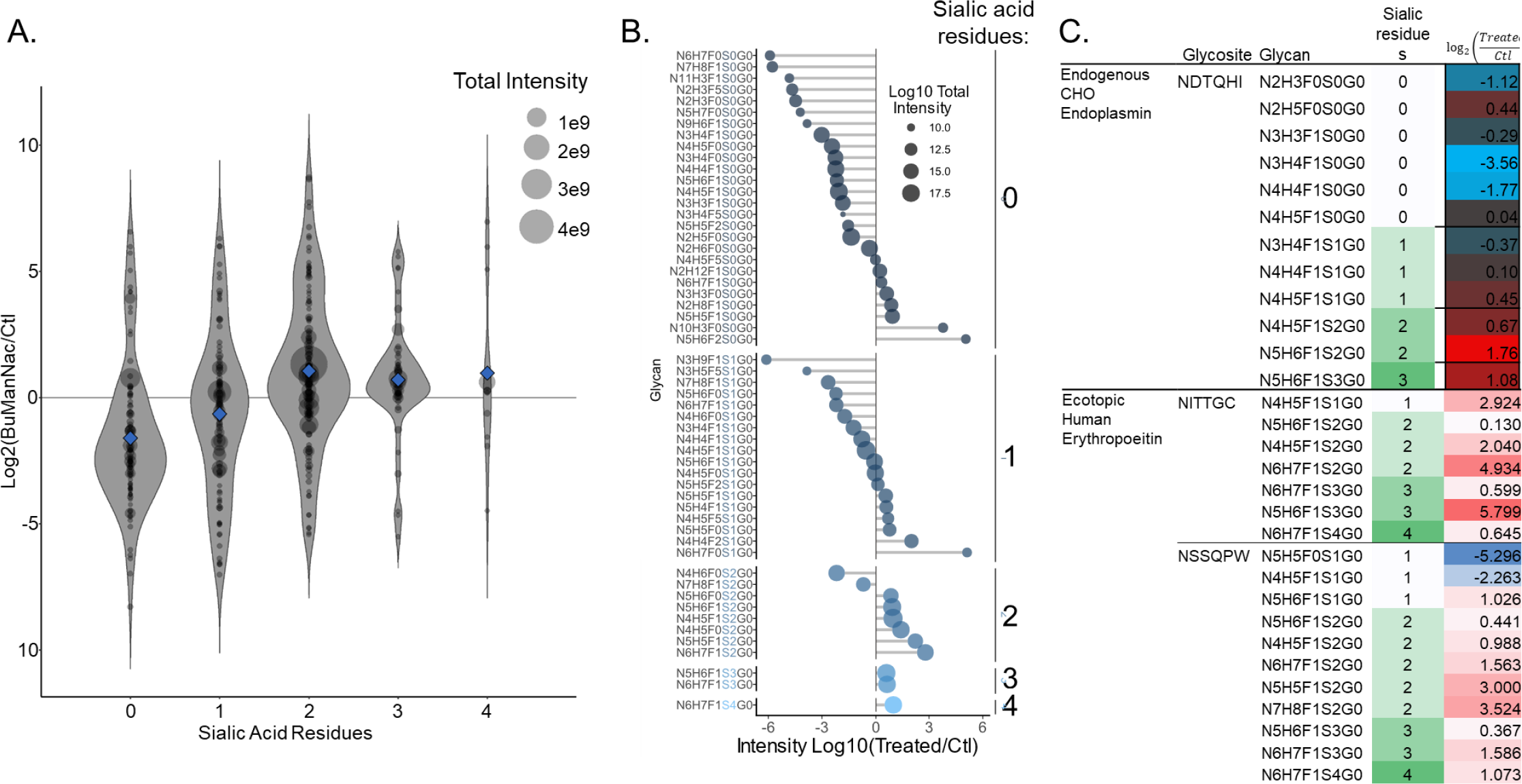
Treatment with the sugar analog 1,3,4-O-Bu_3_ManNAc caused a strong shift toward increased sialic acid residues within identified glycopeptides, including the ectopically expressed human erythropoeitin. A) The overall intensity of glycopeptides containing zero or one sialic acid residues dropped in samples treated with 1,3,4-O-Bu_3_ManNAc with a concomitant increase in intensity for those containing two to four residues. B) Overall intensity changes versus control for all unique identified glycans reveals almost all glycans containing zero sialic residues decreased in intensity, with similar but slightly weaker trend for glycans with 1 sialic residue, while almost all containing 2 or more residues increased. C) Glycosites on two representative proteins bear out the overall trend, with the NDTQHI glycosite on endogenous endoplasmin and the NITTGC and NSSQPW glycosites on ectopic human erythropoietin exhibiting intensity decreases in glycans with 0 or 1 sialic residue and increases in those with 2-4 sialic residues.

More focused analysis of the glycans on particular proteins yielded similar results. Of particular importance to exogenous protein production, the primary utility of CHO cell culture, the increase in sialylation was also observed in the human erythropoietin protein expressed by these cells (Fig. 7C). Consistent with the fact that 1-3-4-O-Bu_3_ManNAc did not increase the number of glycosylation sites system-wide, the same two glycosites on this protein were observed in control and 1-3-4-O-Bu3ManNAc samples. However, both sites exhibited glycan structures with significantly more sialic acid residues.

## Discussion

The increased and largely non-overlapping *glycosite* coverage across the proteome versus SPEG and HILIC in combination with its capacity for *intact glycopeptide* analysis proves NGAG as a valuable glycoproteomics workflow. Overall, we identified 953 unique glycosites when combining all three methods and NGAG contributed to 61% of these, HILIC 50%, and SPEG 32%. In our hands, SPEG was the weakest overall performer with only 301 total glycosite identifications and only 63 that were not observed in other workflows, HILIC identified 477 (a 58% increase over SPEG) and 192 not observed in other workflows, and NGAG identified 586 total glycosites (a 95% increase over SPEG and 23% increase over HILIC) including 386 not seen in the other workflows. Strikingly, only 200 of the total 586 (34%) identifications from NGAG were also observed in the other workflows, leaving a large majority of 386 - or more than the entire SPEG dataset - unidentified by either of the other two. Meanwhile SPEG and HILIC overlapped strongly with each other, with 70% of SPEG glycosites identified by HILIC and 44% vice versa. This large portion of the NGAG identifications that were not observed in either of the two other workflows suggests NGAG removes some enrichment bias or is biased in a different way than SPEG and HILIC. This can most likely be attributed to the initial immobilization step of covalent binding between peptide amino termini and aldehyde resin during initial immobilization, as opposed to immobilization via cis carboxyls (SPEG) or polar nature of glycans (HILIC). Indeed, in theory, virtually all tryptic peptides across the proteome possess α-aminos and should bind the aldehyde solid phase. Highly efficient and robust chemical and enzymatic modifications by guanidine, aniline, PNGaseF, and Asp-N combine to produce a reliable, thorough, and bias-free workflow. The increase in sialylation observed throughout the proteome- and perhaps most importantly, the exogenously expressed EPO – suggest 1-3-4-O-Bu3ManNAc to be a viable glycoengineering tool for improved glycosylation patterns in therapeutic protein production.

## Acknowledgements

The authors would like to thank members of the Zack and Zhang laboratories and Dr. Christina Nemeth for their valuable feedback.

